# Sex differences in the neural circuitry of aggression

**DOI:** 10.64898/2025.12.18.695285

**Authors:** Susan D. Lee, Dario Aspesi, Kim L. Huhman, H. Elliott Albers

## Abstract

The social behavior neural network (SBNN) is a circuit composed of reciprocally connected limbic structures that regulate a range of social behaviors, including aggression. Although both males and females of many species display aggressive behavior, studies of the neural circuitry underlying aggression have focused almost exclusively on males. In the present study, we investigated sex differences in neuronal activation of the neural circuitry controlling aggression in Syrian hamsters (*Mesocricetus auratus*). We employed c-Fos immunohistochemistry to quantify neuronal activation following aggressive encounters between same-sex male and female dyads. Animals were tested in their home cage either alone (n=7 per sex) or with a same-sex, non-aggressive intruder (n=7 per sex) for 10 minutes. Our data revealed substantial sex differences in the neuronal activation of the SBNN following aggression. In some regions, neuronal activity changed in *opposite* directions in males and females compared to controls (e.g., posterior lateral septum), while in others, there was a change in neuronal activation in only one sex (e.g., medial amygdala). These findings support the hypothesis that the neural circuitry regulating aggression exhibits marked sexual differentiation.

**Highlights:**

- The neural circuitry regulating aggression was differentially activated in males and females following aggressive encounters
- Several brain regions exhibited opposite patterns of activation in males and females following aggression
- Only the BNST had the same pattern of changes in neural activity in males and females following aggression

## 1. Introduction

Aggression is exhibited by virtually all vertebrate species, including birds, fish, rodents, and primates, to ensure survival and reproductive success. Aggressive behavior occurs in a variety of social situations, such as during competition for territory, resources, or mates, and it plays a key role in social status and the formation of dominance relationships in both males and females. Offensive aggression refers to proactive attacks frequently aimed at establishing or maintaining dominance, as opposed to defensive aggression, which occurs in response to a threat (Neumann et al., 2010). Despite the widespread occurrence of aggression, most studies investigating the neural mechanisms of aggression have focused almost exclusively on inter-male aggression (Delville et al., 1996, 2000; Lischinsky & Lin, 2020; Veenema & Neumann, 2007; Zhu et al., 2024). This is partly because females of many commonly employed laboratory rodents, such as many strains of mice and rats, do not display the same types of aggression as do males, making direct comparisons difficult. As a result, the neural basis of female aggression remains relatively poorly understood although female aggression has recently been studied more frequently (Been et al., 2019; Oliveira & Bakker, 2022; Pandolfi et al., 2021). Syrian hamsters (*Mesocricetus auratus*) have proven to be a valuable model because both males and females readily display high levels of offensive aggression and readily form hierarchical dominance relationships, allowing for direct study of sex differences.

The social behavior neural network (SBNN) is a neural circuit composed of reciprocally connected brain structures thought to regulate a variety of social behaviors, including aggression (Albers, 2012, 2015; Caldwell, 2017; Goodson & Kingsbury, 2013; Newman, 1999; O’Connell & Hofmann, 2011, 2012). Over the years, numerous studies have examined elements of the neural circuitry of aggression in a variety of species, including in male hamsters (Delville et al., 2000). In these studies, limbic structures such as the ventromedial hypothalamus (VMH), amygdala, lateral septum (LS), bed nucleus of the stria terminalis (BNST), and ventrolateral hypothalamus were found to be activated following aggressive encounters (Delville et al., 2000).

Although both male and female hamsters, like many primates, readily display offensive aggression and form hierarchical dominance relationships, it is not known whether the underlying neural mechanisms are the same in both sexes (Albers et al., 2002). Recent evidence suggests that there are substantial sex differences in the neural mechanisms regulating aggression and social status in hamsters. For example, in the anterior hypothalamus (AH), serotonin (5-HT) and vasopressin (AVP) act in opposite ways to modulate the expression of aggression and dominance in males and females (Terranova et al., 2016). In addition, acquisition of dominance is associated with activation of 5-HT neurons within the dorsal raphe (DR) in females but activation of hypothalamic AVP neurons in males (Terranova et al., 2016).

In the present study, we directly compared neural activity within the aggression-regulating circuitry in male and female Syrian hamsters. Using c-Fos immunohistochemistry as a marker of neuronal activation, we quantified activity in key SBNN regions following aggressive encounters to test the hypothesis that sex differences exist in the neural mechanisms underlying offensive aggression. Understanding these differences will advance our knowledge of the neural regulation of social behavior and may provide insight into sex-specific vulnerabilities in aggression-related neuropsychiatric disorders.

## 2. Methods

### 2.1 Subjects

Adult male and female Syrian hamsters (*Mesocricetus auratus)* were bred in the Georgia State University vivarium or obtained from Charles River Laboratories Inc. (Wilmington, MA). Animals were housed under a 14:10 light:dark cycle as is customary in hamsters to maintain gonadal patency. Adult male and female hamsters, weighing at least 120g, were individually housed in 24 cm X 33 cm X 20 cm polycarbonate cages filled with corncob bedding and nesting material for four weeks before behavioral testing to allow residents to establish home cage territory. It is important to note that individual housing is not stressful for Syrian hamsters (Ross et al., 2017). All hamsters had *ad libitum* access to food and water. The estrous cycle of female hamsters was monitored daily by vaginal discharge beginning 8 days before the start of behavioral testing. Males were handled daily as a yoked control for handling females. The four-day estrous cycle contains the following days: diestrus 1 (D1), diestrus 2 (D2), proestrus (P), and estrus (E). Females underwent behavioral testing on D1 only, with males yoked to each female to ensure that a similar number of males and females were tested each day. All animals were tested in the first three hours of the dark phase of the daily light-dark cycle to minimize circadian variation. Stimulus hamsters used as non-aggressive intruders (NAI) were younger, smaller (100-110g), and group-housed to ensure that the resident subjects would exhibit aggression. All procedures and protocols were performed in accordance with the principles of the National Institutes of Health Guide for the Care and Use of Laboratory Animals and were approved by the Institutional Animal Care and Use Committee at Georgia State University.

### 2.2 Behavioral Testing

Male and female hamsters were observed for agonistic behavior during a 10-min encounter with a same-sex NAI. Subjects remained in their home cage, and a novel same-sex NAI was introduced (n=7 per sex). All residents displayed aggressive behaviors towards the NAI. Hamsters in the no-social control group (n=7 per sex) were transported to the behavioral testing suite on test day but remained alone in their home cage. Test sessions were recorded, and videos were scored using Noldus Observer (11.5, Leesburg, VA) by observer blind to the treatment group. Inter-rater reliability, or percent agreement between observers, was above 90%. The total duration of the four classes of behavior (aggression, submission, social, and non-social behavior, as described in detail in (Albers et al., 2002)) were quantified. Aggression included pins, upright and side attacks, bites, chases, and anogenital sniffing. Submissive, or defensive, behavior included flight, avoidance, tail up, upright stance with paws up, stretch attend, head flag, and attempted escape from the cage. Social behavior included stretch, approach, sniffing, nose touching, and flank marking. Non-social behavior included grooming, locomotion, exploration, nesting, feeding, and sleeping.

### 2.3 Transcardial perfusions

Ninety minutes after testing, animals were deeply anesthetized with sodium pentobarbital and perfused transcardially. An incision was made into the chest to expose the heart. A needle connected to surgical tubing was inserted into the left ventricle, and the right atrium was pierced with surgical scissors. The tubing was run through a peristaltic pump set to 15 ml/minute. Animals were first perfused with 100 ml cold phosphate buffer solution (PBS) and then with 100 ml cold 4 % paraformaldehyde (PFA) in PBS. Brains were extracted, post-fixed in 4 % PFA for 24h and then placed in 30 % sucrose in PBS for cryoprotection until sectioning.

### 2.4 c-Fos immunohistochemistry

Brains were sectioned coronally at 40μm on a cryostat, and sections were stored in a cryoprotectant solution (500 mL PBS, 300 g sucrose, 10 g polyvinyl pyrrolidone, 300 mL ethylene glycol) at −20°C until c-Fos immunohistochemistry was performed. For each region of interest (ROI), three to four sections per animal were isolated and rinsed five times with PBS to remove excess cryoprotectant. Sections were blocked in 10 % normal donkey serum (NDS) in PBS with 1 % Triton-X for 1hr. Then sections were incubated in primary antibody (ab208942 mouse anti-c-Fos, 1:500 dilution, Abcam, Boston, MA) diluted in 5 % NDS in PBS with 1 % Triton-X overnight on a shaker at room temperature. Sections were rinsed 3 x 5 min in PBS and incubated for 2hr at room temperature in secondary antibody (Alexa Fluor 594 conjugated donkey anti-mouse IgG, 1:250 dilution, Jackson ImmunoResearch Laboratories, West Grove, PA). Sections were mounted onto Super Frost Plus slides and coverslipped with Vectashield Hard Set Mounting Medium for Fluorescence with DAPI (H1500, Vector Laboratories, Burlingame, CA). Images were collected on the Keyence microscope at 4x magnification. Using the Syrian hamster stereotaxic atlas (Morin & Wood, 2001), ROI templates for each brain region were generated from DAPI-stained sections, and these templates overlayed each section during cell quantification to ensure that the same size area was counted across animals per region (Supplemental Fig. 1). Quantification of c-Fos-positive cells was performed with the cell counter plugin on FIJI. The regions quantified include the hypothalamus ((paraventricular nucleus (PVN, ROI: 0.555 mm^2^), medial supraoptic nucleus (mSON, ROI: 0.606 mm^2^), anterior hypothalamus (AH, ROI: 0.947 mm^2^), ventromedial hypothalamus (VMH, ROI: 0.478 mm^2^), lateral hypothalamus (LH, ROI: 1.11 mm^2^)), amygdala ((medial amygdala (MeA, ROI: 1.237 mm^2^), central amygdala (CeA, ROI: 0.536 mm^2^), basolateral amygdala (BLA, ROI: 0.942 mm^2^)), bed nucleus of the stria terminalis (BNST, ROI: 1.823 mm^2^), lateral septum (LS, see Fig. 1 legend for ROI area sizes), periaqueductal grey (PAG, ROI: 4.638 mm^2^), dorsal raphe (DR, ROI: 2.150 mm^2^), and the preoptic area (POA, ROI: 2.469 mm^2^). Given the distinct anatomical and functional differences of the LS, anterior and posterior LS were analyzed separately (Fig. 1). Brain regions were quantified bilaterally in 2-4 sections per brain region and averaged for each subject.

**Figure 1.**
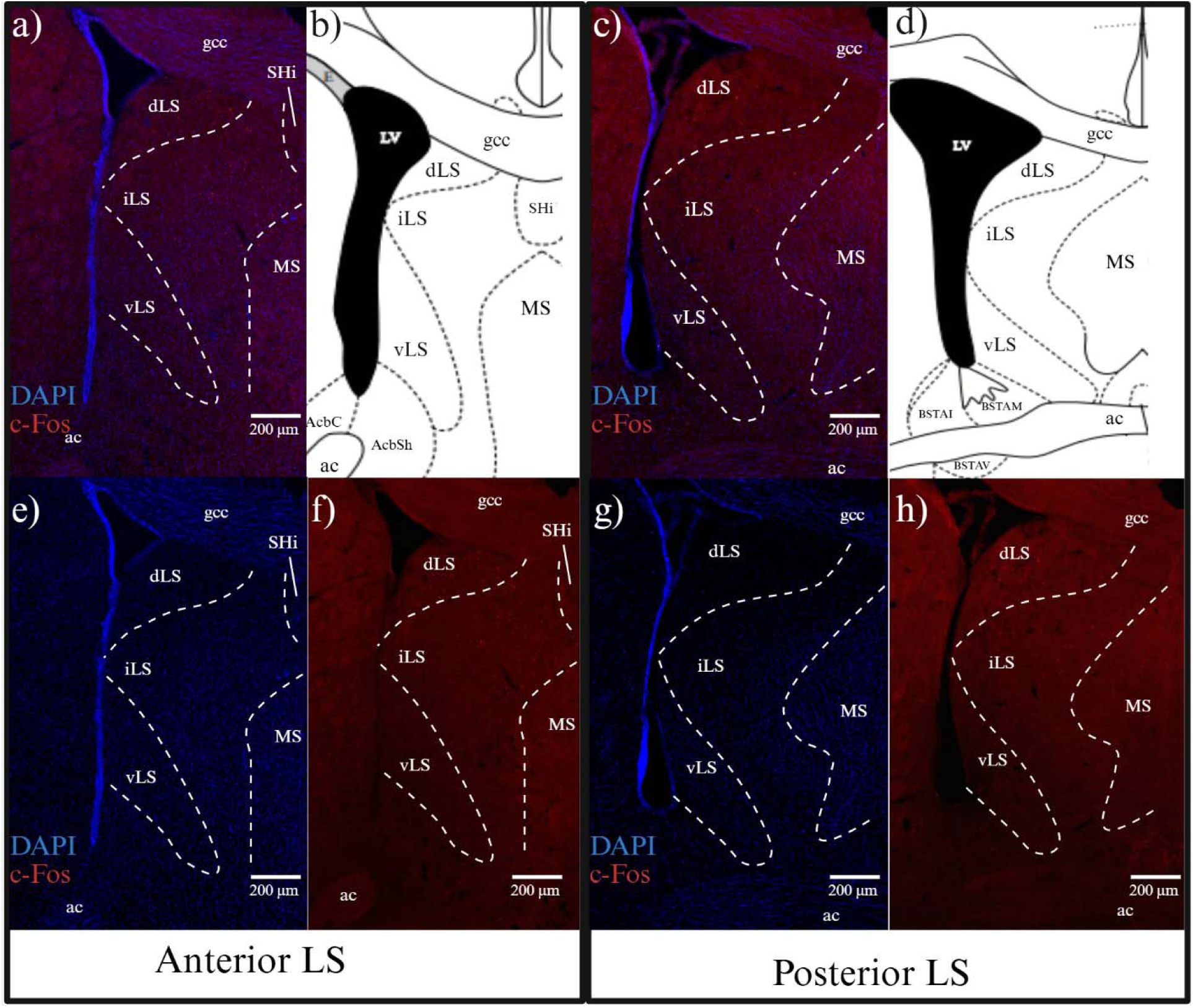
Representative images of anterior and posterior lateral septum (LS) subregions at 4X magnification. a,e,f) Representative image of the dorsal (ROI: 0.482 mm^2^), intermediate (ROI: 0.359 mm^2^), and ventral (ROI: 0.767 mm^2^) subregions of the anterior LS. b,d) Schematic illustration depicts the dorsal (ROI: 0.721 mm^2^), intermediate (ROI:1.023 mm^2^), and ventral (ROI:0.892 mm^2^) subregions of the anterior and posterior LS at +1.55 mm (anterior) and +0.8 mm (posterior) from Bregma (adapted from Morin and Wood 2001). c,g,h) Representative image of the dorsal, intermediate, and ventral subregions of the posterior LS with DAPI (blue) and c-Fos (red). dLS=dorsal LS, iLS = intermediate LS, vLS = ventral LS, MS = medial septum, ac = anterior commissure, gcc=genu of the corpus collosum, SHi = septohippocampal nucleus, AcbSh = nucleus accumbens shell, AchC = nucleus accumbens core, BSTAM = anteromedial part of the bed nucleus of the stria terminalis (BNST), BSTAI = anterointermediate part of the BNST, BSTAV = anteroventral part of the BNST, LV = lateral ventricle.

### 2.5 Statistical analysis

Data were analyzed in SPSS version 29 (IBM). Graphs, generated in GraphPad PRISM, are of original data and not transformed data. Normality (Shapiro-Wilk) and homogeneity of variance (Levene’s test) were assessed before analysis. Two-way between-subjects analysis of variance (ANOVA) with Bonferroni’s post-hoc test were used to examine group differences when assumptions were met. If the data violated the assumptions for ANOVA, then the non-parametric test Mann-Whitney U was used. Because non-parametric factorial tests are limited, these analyses focus on simple effects rather than formal interactions. Statistical significance was conferred at p<0.05. Outliers (≥2 SD from the mean) were excluded from statistical comparisons: one male (social group) for anterior LS and POA, and one female (control group) for VMH. Effect sizes were reported as partial eta squared (η ^2^) for parametric tests and eta squared (η^2^) for non-parametric tests.

## 3. Results

### 3.1 Behavior analysis

While both males and females displayed aggression, females engaged in significantly more aggressive behavior than did males (t(12)=6.637, p<0.001, η ^2^=0.77; Fig.2). Conversely, males spent more time in non-social behavior than did females (t(12)=-7.424, p<0.001, η ^2^=0.81). No sex differences were observed in social behavior (U=37.00, z=1.597, p=0.110). Neither male nor female hamsters engaged in submissive behavior.

**Figure 2.**
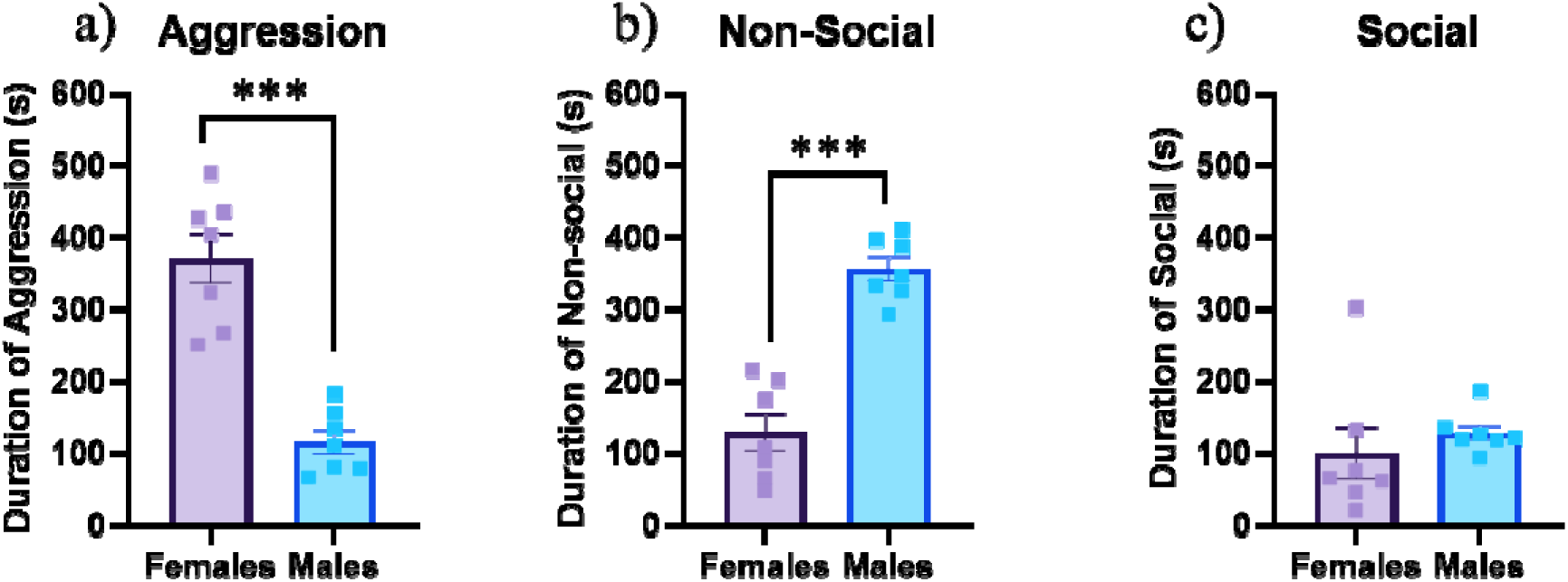
Duration of behaviors: aggression (a), non-social behavior (b), and social behavior (c). No submissive behaviors were observed during behavioral testing. The data are expressed as means ± SEM with individual data points shown as square. ***p<0.001

### 3.2 Quantification of c-Fos immunoreactivity (ir)

The number of c-Fos-ir (+) cells was quantified 90 minutes following control or aggression test sessions in the brain regions indicated in Tables 1 and 2. In the anterior LS (antLS), there was no significant main effect of sex (F_1,24_=0.057, p=0.813) or group (F_1,24_ = 0.841, p=0.368), nor a significant interaction (F_1,24_=2.317, p=0.141; Fig. 3a). In contrast, the posterior LS (postLS) showed a significant sex x group interaction with a large effect size (F_1,24_=10.083, p=0.004, η_ρ_^2^= 0.296). Post-hoc analysis confirmed that males had greater c-Fos-ir compared to females (p=0.008) following aggression. Post-hoc analysis also showed a significant difference in c-Fos-ir between control and aggression groups within each sex: c-Fos expression was lower in females following aggression (p=0.032) but higher in males following aggression (p=0.037) compared to their control counterparts (Fig. 3b,4).

**Figure 3.**
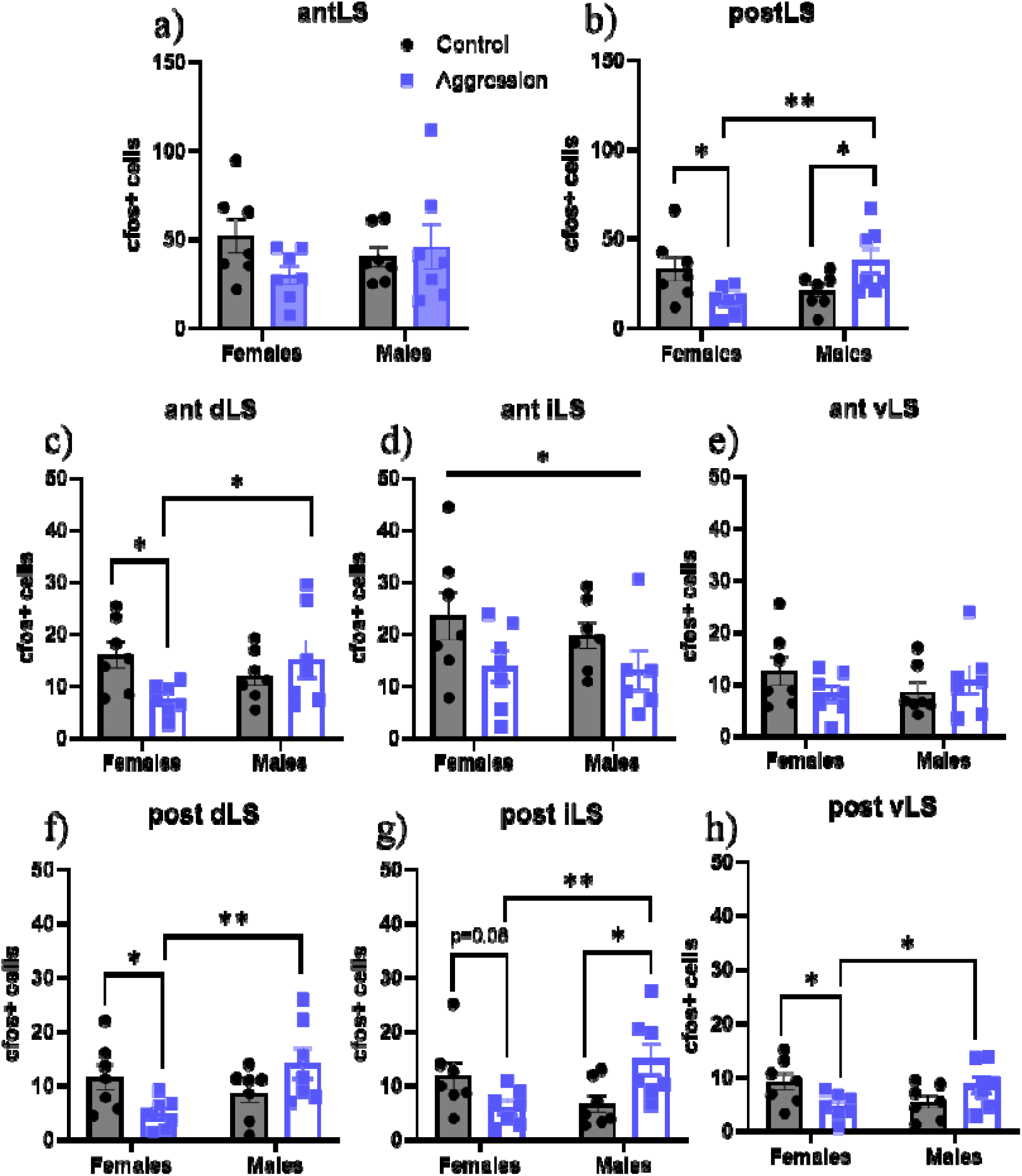
Neural activity in the lateral septum (LS) measured with c-Fos immunoreactivity (ir). a) No significant differences in neural activity were observed in the antLS. b) c-Fos-ir was lower in females but higher in males following aggression compared to controls in the postLS. Males have more c-Fos-ir after aggression compared to females. c) c-Fos-ir was lower in females following aggression in ant dLS compared to controls. Males had higher c-Fos-ir after aggression compared to females. d) Both males and females have lower c-Fos-ir after aggression in the ant iLS compared to controls e) No significant differences were observed in the ant vLS. f) c-Fos-ir was lower in females following aggression in the post dLS compared to controls. Males had c-Fos-ir after aggression in males compared to females. g) c-Fos-ir tends to decrease in females, but increases in males following aggression in the post iLS. Males show more c-Fos-ir following aggression compared to females. h) c-Fos-ir was lower in females following aggression compared to controls. Males had more c-Fos-ir following aggression compared to females. Error bars indicate standard error of the mean. *p<0.05, **p<0.01, #p<0.09 with medium-large effect size. Regions were defined using templates to ensure the same area was counted across animals.

**Figure 4.**
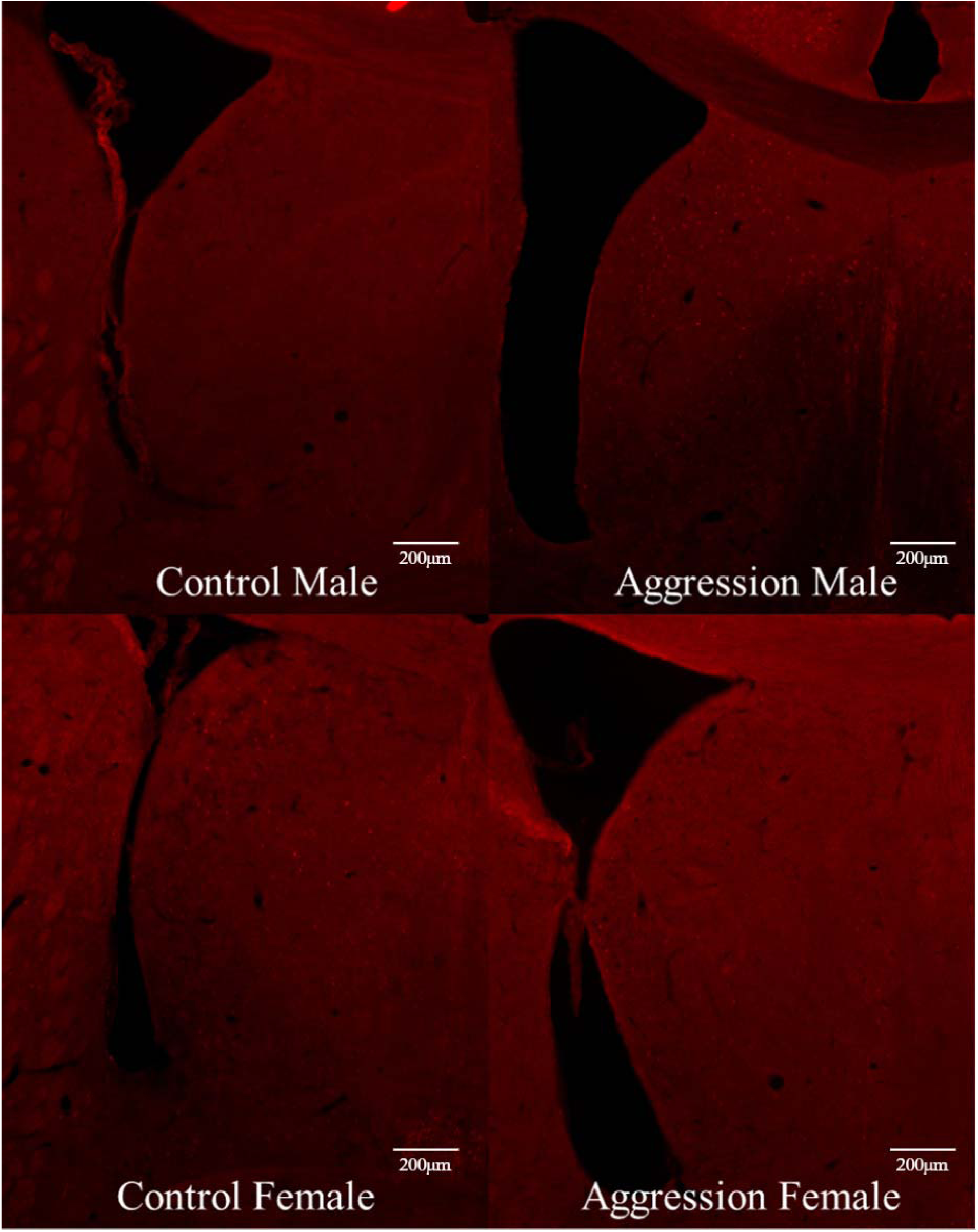
Representative images of the LS at 4X showing differences in c-Fos-ir cells between control and aggression groups in females and males.

**Table 1.** Summary of changes in c-fos-immunoreactivity (ir) within sex following aggression compared to no-social control condition, as well as differences in c-fos-ir between sexes. * indicates significance, conferred at p<0.05, # indicates trending to significance with a medium-large effect, conferred at p<0.09.

| Brain Regions | Females | Males | No-Social Control | Aggression |
| --- | --- | --- | --- | --- |
| Lateral Septum |  |  |  |  |
| Anterior LS | - | - | - | - |
| Posterior LS | Lower (*) | Higher (*) | - | M>F (*) |
| BNST | Higher (*) | Higher (#) | - | F>M (*) |
| MPOA | - | - | F>M (*) | - |
| Hypothalamus |  |  |  |  |
| mSON | Lower (#) | Higher (*) | - | M>F (*) |
| PVN | - | - | - | - |
| AH | - | - | - | - |
| LH | Lower (*) | - | - | M>F(*) |
| VMH | Higher (*) | - | - | - |
| Amygdala |  |  |  |  |
| MeA | - | Higher (*) | - | - |
| CeA | - | - | - | - |
| BLA | - | - | - | - |
| DR | Lower (*) | Higher (#) | F>M (*) | - |
| PAG | - | - | - | - |

**Table 2.** Summary of changes in c-Fos-immunoreactivity (ir) in the subregions of the lateral septum (LS) within sex following aggression compared to the no-social control condition. * indicates significance, conferred at p<0.05, # indicates trending to significance with a medium-large effect, conferred at p<0.09.

| **Brain Regions** | **Females** | **Males** | **F v M** |
| --- | --- | --- | --- |
| Lateral Septum |  |  |  |
| *Anterior LS* | - | - | - |
| Ant dLS | Lower (*) | - | No difference during control condition<br>M>F following aggression(*) |
| Ant iLS | Lower (*) | Lower (*) | - |
| Ant vLS | - | - | - |
| *Posterior LS* | Lower (*) | Higher (*) | No difference during control condition<br>M>F following aggression(*) |
| Post dLS | Lower (*) | - | No difference during control condition<br>M>F following aggression(*) |
| Post iLS | Lower (#) | Higher (*) | No difference during control condition<br>M>F following aggression(*) |
| Post vLS | Lower (*) | - | No difference during control condition<br>M>F following aggression(*) |

Given the anatomical and functional heterogeneity of the LS, we conducted a subregional analysis of the dorsal, intermediate, and ventral LS (Table 2). In the anterior dorsal LS (ant dLS), a significant sex x group interaction with a large effect size (F_1,24_=5.674, p=0.025, η_ρ_^2^=0.191) was observed such that females had less c-Fos-ir following aggression compared to no-social controls (p=0.023). In males, no differences in c-Fos-ir were observed between the aggression group and the no-social controls. Males expressed more aggression-induced c-Fos-ir than did females (p=0.036; Fig. 3c). In the anterior intermediate LS (ant iLS), a significant main effect of group (F_1,24_=5.039, p=0.035, η_ρ_^2^=0.180) was observed where both males and females had lower c-Fos-ir following aggression (Fig. 3d). There were no significant differences in c-Fos-ir across groups in the anterior ventral LS (ant vLS) (Fig. 3e).

In the posterior dorsal LS (post dLS), c-Fos-ir was lower in females (U=8.00, z=-2.109, p=0.035, η^2^=0.342) following aggression compared to no social controls and to males in the aggression group. No significant differences were observed between groups in males (U=33.00, z=1.086, p=0.277; Fig. 3f). In the posterior intermediate LS (post iLS), a significant sex x group interaction was observed (F_1,24_=10.137, p=0.004, η_ρ_^2^=0.297): males had greater amounts of c-Fos-ir after aggression compared to no-social controls (p=0.013), whereas females tended to have lower amounts of c-Fos-ir with a medium effect size (p=0.081, η_ρ_^2^ = 0.122) following aggression (Fig. 3g). Similarly, in the posterior ventral LS (post vLS), a two-way sex x group interaction revealed significantly lower levels of c-Fos-ir (F_1,24_=7.954, p=0.009, η_ρ_^2^ =0.249), with post-hoc analyses showing lower c-Fos-ir in females following aggression compared to controls but no significant difference in males compared to their control counterparts (p=0.03; Fig. 3h). In the dorsal (p=0.005), intermediate (p=0.009), and ventral (p=0.05) postLS, c-Fos-ir was higher in males compared to females after aggression (Fig. 3f-h).

In the BNST, there was a significant main effect of sex (F_1,24_=9.709, p=0.005, η ^2^ =0.288) and aggression (F_1,24_=12.067, p=0.002, η ^2^ =0.335) but no significant sex x group interaction (F_1,24_=0.582, p=0.453; Fig. 4a). Post-hoc tests showed a significantly greater amount of c-Fos-ir in females (p=0.006) and a trend toward more c-Fos-ir in males (p=0.067, η ^2^ =0.133) following aggression compared to controls. A sex difference was observed where females had greater c-Fos-ir following aggression compared to males (p=0.011).

In the MPOA, a significant main effect of sex (F_1,23_=12.703, p=0.002, η ^2^ =0.356) was observed but not a significant main effect of aggression (F_1,23_=0.044, p=0.836) nor a significant interaction (F_1,23_=1.291, p=0.268; Fig. 4b). The main effect of sex was driven by greater c-Fos-ir in control females compared to control males in the MPOA (p=0.003).

Sex differences in c-Fos-ir in the aggression group were observed in the hypothalamus. In the mSON, a significant sex x group interaction (F_1,24_=8.646, p=0.007, η ^2^ =0.265) was observed (Fig. 4c). Post-hoc analysis revealed significantly greater c-Fos-ir in males in the aggression group than in no-social controls (p=0.037) and compared to females in the aggression group (p=0.011). A trend toward lower c-Fos-ir in females (p=0.063), with a large effect size (η ^2^ = 0.136), was also observed in the mSON. In the LH, females expressed less c-Fos-ir following aggression compared to no-social controls (U=7.50, z=-2.175, p=0.030, η^2^ =0.364) while males showed no change (U=28.00, z=0.447, p=0.655). Males expressed more c-Fos-ir than their female counterparts following aggression (U=43.50, z=2.430, p=0.015, η^2^ =0.454; Fig. 4d). In the PVN, there was no significant main effect of sex (F_1,24_=0.869, p=0.361) or group (F_1,24_=0.325, p=0.574), nor a significant interaction (F_1,24_=1.213, p=0.282; Fig. 4e). In the VMH, c-Fos-ir increased in females following aggression compared to their control counterparts (U=39.00, z=2.582, p=0.01, η^2^ =0.513), but not in males (U=33.50, z=1.151, p=0.25; Fig. 4f). In the AH, no significant main effects of sex (F_1,24_=0.103, p=0.751) or group (F_1,24_=0.139, p=0.712) were observed, although there was a trend toward a significant sex x group interaction (F_1,24_=3.503, p=0.074; η ^2^ = 0.127; Fig. 4g).

In the MeA, males showed greater c-Fos-ir after aggression compared to no-social controls (U=43.00, z=2.364, p=0.018, η^2^ =0.430), whereas females did not (U=35.00, z=1.342, p=0.180; Fig. 4h). No sex differences were observed in either control or aggression conditions. In the CeA and BLA, c-Fos-ir did not differ between groups or sexes (Fig. 4i-j).

In the PAG, no significant main effects of sex (F_1,24_=0.560, p=0.462) or group (F_1,24_=0.009, p=0.923), nor a significant interaction (F_1,24_=1.508, p=0.231), were observed (Fig. 4k). In the DR, c-Fos-ir was lower in females following aggression compared to no-social controls (U=7.50, z=-2.175, p=0.03, η^2^ =0.364) and there was a trend for greater c-Fos-ir in males (U=34.00, z=1.857, p=0.063; η^2^ = 0.287; Fig. 4l). Control males also expressed less c-Fos-ir compared to control females (U=1.00, z=-2.857, p=0.004, η^2^ =0.628).

## 4. Discussion

The present study provides strong support for the hypothesis that the neural circuitry regulating aggression in Syrian hamsters is sexually differentiated. We observed significant sex differences in c-Fos-ir in a number of key structures of the SBNN between the groups that expressed aggression and sex-matched controls (Fig. 5, Table 1). Notable, some of the sex differences in neuronal activation within the SBNN may reflect the sex differences in the amount of aggression displayed. While both sexes engaged in aggressive behavior, females were significantly more aggressive than males. Across the 14 brain regions quantified, males and females showed changes in the *same* direction following aggression only in the posterior BNST. Of note, several regions of the SBNN exhibited changes in *opposite* directions in neuronal activation between sexes, with a significant interaction of sex and aggression observed in the postLS and mSON, and a similar trending pattern in the DR. Other regions, such as the MeA, LH, and VMH, showed significant changes in neuronal activation in only one sex. Together, these findings indicate that the sex differences in neural activation after aggression are robust and reflect genuine sexual variation in the functioning of the SBNN, rather than merely differences in the amount of aggressive behavior.

**Figure 5.**
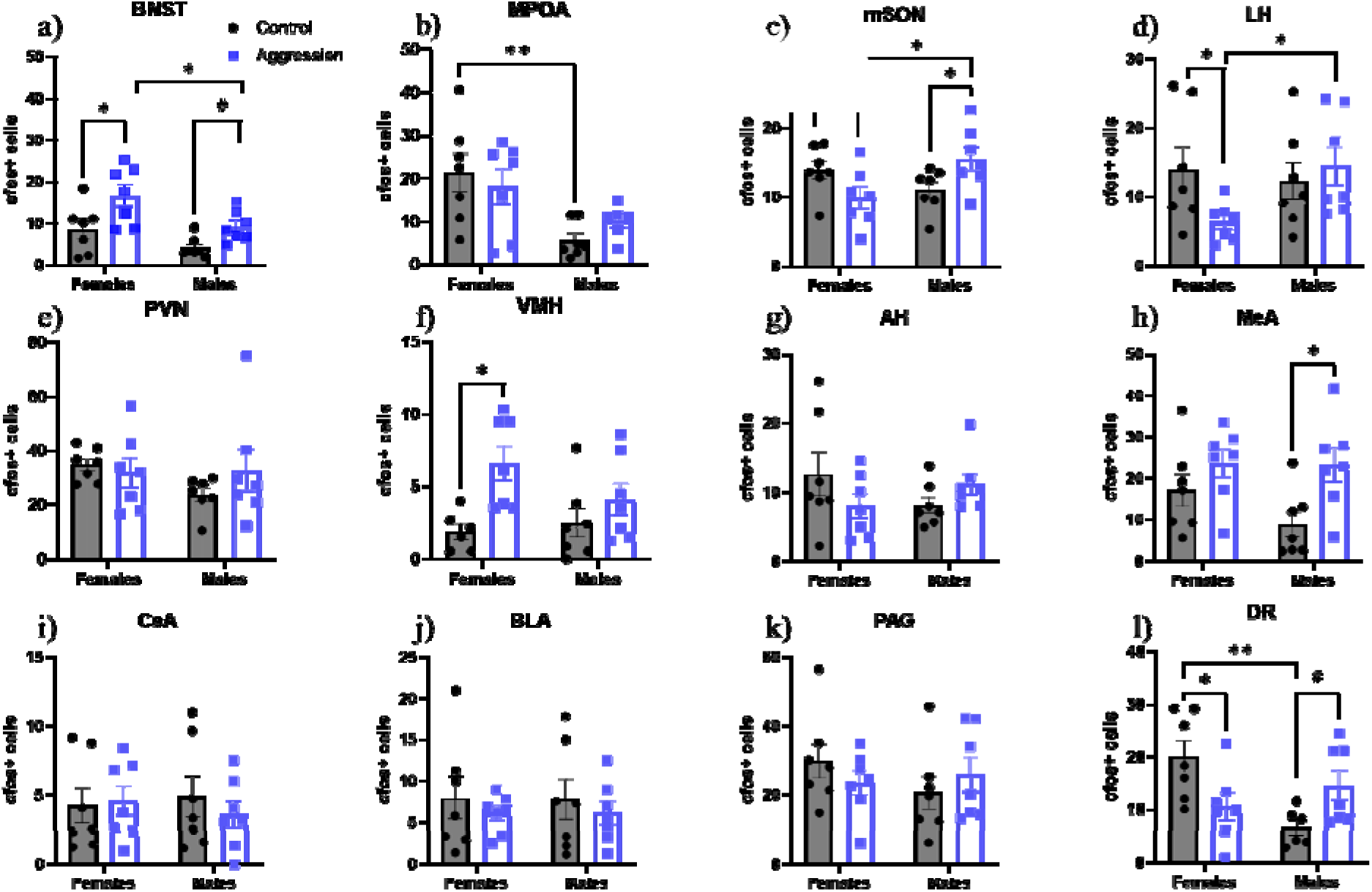
Neural activity in the SBNN measured with c-Fos immunoreactivity (ir). a) c-Fos-ir was higher in both males and females following aggression in the BNST than in controls. Females had more c-Fos-ir following aggression compared to males. b) Females had more c-Fos-ir compared to males in the control condition in the MPOA. c) There was a trend for c-Fos-ir to be lower in females following aggression compared to controls but c-Fos-ir was greater in males following aggression than in controls. Males had more c-Fos-ir following aggression compared to females. d) c-Fos-ir was lower in females following aggression compared to controls. Males have more c-Fos-ir following aggression compared to females. e) No differences in c-Fos-ir were observed in the PVN. f) c-Fos-ir was higher in females following aggression than in controls in the VMH. g) No significant differences in c-Fos-ir were observed in the AH. h) c-Fos-ir was greater in males following aggression than in controls. i-k) No significant differences are observed in the CeA, BLA, or PAG. l) c-Fos-ir was significantly lower in females following aggression compared to controls and significantly greater in males compared to controls. Females have more c-Fos-ir compared to males in the control condition. Error bars indicate standard error of the mean. Individual data points are expressed as black circles for control condition and as blue square for hamsters exposed the aggression condition. *p<0.05, **p<0.01, #p<0.09 with medium-large effect size. Regions were defined using ROIs to ensure the same area was counted across animals.

**Figure 6.**
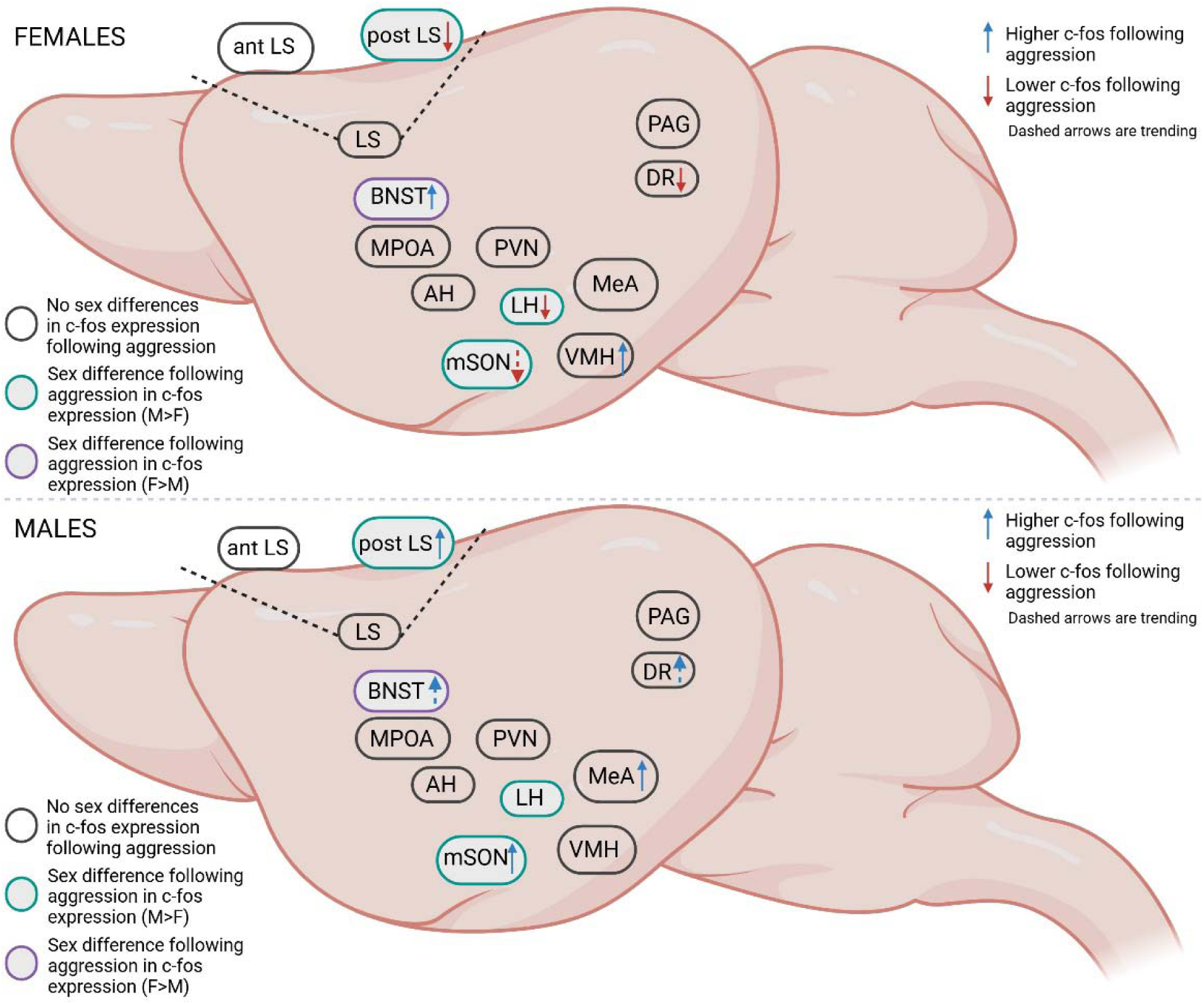
Summary of the neural activity in the SBNN following aggression compared to controls.

The LS has long been known to have an inhibitory role in the regulation of aggression in males, wherein pharmacological, and more recently chemogenetic and optogenetic, inhibition of the LS increases aggression, while activation decreases it (Borland et al., 2020; Leroy et al., 2018; McDonald et al., 2012; Menon et al., 2022; Potegal et al., 1981; Rizzi-Wise & Wang, 2021; Wong et al., 2016). Interestingly, however, Delville et al. found that c-Fos expression measured across the entire LS in male hamsters did not change in response to an aggressive encounter (Delville et al., 2000). Given the anatomical and functional heterogeneity of the LS (Besnard & Leroy, 2022; Menon et al., 2022; Rizzi-Wise & Wang, 2021; Wirtshafter & Wilson, 2021), we examined c-Fos expression across LS subregions along both the dorsal-ventral and rostral-caudal axes. These analyses suggest that: 1) the posterior LS may play a more important role in aggression than the anterior LS, 2) this role may be opposite in males and females, and 3) neuronal activation patterns vary across dorsal, intermediate, and ventral subregions of the LS following aggression.

There is an increasing body of evidence that the LS regulates aggression through different neurochemical signals in a sex-specific manner. In male mice, optogenetic activation of the LS decreased aggression, whereas optogenetic inhibition increased it (Wong et al., 2016), consistent with pharmacological data (Borland et al., 2020). In contrast, AVP signaling in the LS appears to have opposite effects on aggression across sexes. In male rats, AVP release in the LS was positively correlated with aggressive behavior (Beiderbeck et al., 2007; Veenema et al., 2010). Strains of high-aggressive male rats displayed elevated levels of AVP release in the LS that predicts shorter attack latencies and increased aggression (Veenema et al., 2010). However, in isolated female rats and female rats trained for aggression, AVP injections in the LS reduced aggression (Oliveira et al., 2021). Interestingly, reduced release of AVP in the dorsal LS combined with increased release of OXT in the ventral LS appears to be required for aggression in these isolated and trained female rats. Similar sex-dependent effects have been observed in social play, a precursor to adult aggression: blocking AVP 1a receptors increased social play in males but decreased it in females (Veenema et al., 2013). In female California mice, greater c-Fos expression in the ventral LS was also linked to higher levels of aggression during diestrus compared to either the proestrous or estrous phase of their estrous cycle (Davis & Marler, 2004). Taken together, these findings highlight the LS as a central hub in the neural circuitry regulating aggression, with its function shaped by sex, neurochemical context, and subregional specialization.

Another key element of the aggression circuitry is the serotoninergic projection from the raphe nuclei. The powerful inhibitory actions of 5-HT on aggression are well established across diverse species (Chien et al., 2024; Muroi & Ishii, 2019; J. C. Nordman et al., 2020; J. Nordman & Li, 2020; Takahashi et al., 2010, 2015; Takahashi, Quadros, et al., 2012; Takahashi, Schilit, et al., 2012), although almost all of these studies were done in males only. For example, inhibition of DR neurons via GABA_B_ receptor activation increased aggression in male mice, an effect dependent on 5-HT receptor activity (Takahashi et al., 2010). More recent studies, however, demonstrate substantial sex differences in serotonergic regulation of aggression. In male hamsters, administration of 5-HT agonists into the hypothalamus or systemic treatment with selective 5-HT reuptake inhibitors suppressed aggression, whereas the same manipulations facilitated aggression in females (Terranova et al., 2016). Our findings extend this model by showing that neuronal activation in the DR is higher following aggression in males but lower in females. These results provide further evidence that specific nodes of the aggression circuitry can operate in an opposite way across sexes.

It is also noteworthy that neuronal activation appeared to change in opposite directions in males and females following aggression in another key site in the aggression circuitry: the mSON. Consistent with previous findings in males (Delville et al., 2000), the present study found greater neuronal activation in the mSON of males following aggression compared to controls. In contrast, females exhibited lower levels of c-Fos-ir in the mSON following aggression than controls. In addition, following aggression, neuronal activation was higher in males than in females. Neurons in the mSON are a major source of AVP innervation to hypothalamic nuclei and other downstream regions, where AVP receptor activation has opposite effects on social communication and aggression in males versus females (Brownstein et al., 1980; De Vries & Miller, 1999; Delville et al., 1994; Ferris et al., 1990; Terranova et al., 2016; Young & Gainer, 2003). The mSON is also a sexually dimorphic area, with greater AVP-ir cells in the mSON of male hamsters compared to females. This anatomical sexual dimorphism is consistent with our functional findings: males showed higher c-Fos expression following aggression, whereas females showed a trend toward decreased activation. Notably, the mSON has bi-directional connections with the LS (Ferris et al., 1990), where we observed a similar sex-specific pattern of change in neuronal activation.

In addition to these pronounced sex differences, where males and females showed opposite patterns of neuronal activation, more nuanced sex differences were also observed in other regions of the circuitry. For example, both sexes exhibited higher c-Fos expression in the BNST in the aggression groups, but the magnitude of this increase was greater in females than in males. A similar increase in neuronal activation in the posterior BNST was previously reported in male hamsters (Delville et al., 2000), as well as in male and female California mice (Davis & Marler, 2004) and Mongolian gerbils (Pan, Zhu, Wang, et al., 2020; Pan, Zhu, Xu, et al., 2020), following an aggressive encounter. In male California mice, greater pERK(+) expression in the posterior BNST was also positively correlated with the number of bites during aggression (Trainor et al., 2010). Together, these findings suggest that, unlike other regions of the SBNN that exhibit strong sex specificity, the BNST may represent a shared hub for the regulation of aggression in both sexes.

In the present study, we observed significantly greater neuronal activation in the VMH of females, but not males, following aggressive encounters. This pattern mirrors previous findings in California mice (Davis & Marler, 2004; Y. Hashikawa et al., 2017; Wong et al., 2016; Yang et al., 2013) and Mongolian gerbils (Pan, Zhu, Wang, et al., 2020; Pan, Zhu, Xu, et al., 2020). For example, female California mice show increased neuronal activation in the VMH after aggression, whereas males do not, as assessed by changes in pERK expression during aggression testing (Trainor et al., 2010). However, evidence from other rodent species indicate that the VMH, particularly the ventrolateral subdivision (VMHvl), is critical for male aggression (Falkner et al., 2016; K. Hashikawa et al., 2017; Y. Hashikawa et al., 2017; Khodai & Luckman, 2021; Wong et al., 2016; Yang et al., 2013; Zhu et al., 2024). Notably, both female Syrian hamsters and female California mice readily engage in aggression, apart from maternal aggression. Additionally, we observed female Syrian hamsters engaged in aggression longer than did the male hamsters. The difference in time spent engaging in aggressive behavior may explain this discrepancy, with the more aggressive females showing higher c-Fos expression in the VMH after aggression compared to their male counterparts.

No significant differences in neuronal activation following aggression were observed in several other sites thought to be part of aggression circuitry including the antLS, MPOA, PVN, CeA, BLA, and PAG in either sex. The lack of robust AH activation in males or females following an aggressive encounter is consistent with prior data showing no significant differences in c-Fos expression in the AH following an aggressive encounter in adult male hamsters (Delville et al., 2000) nor following social play in adolescent males (Cheng et al., 2008). Interestingly, the three regions in which males showed higher c-Fos expression while females had lower c-Fos expression than controls (postLS, mSON, DR) all have reciprocal connections with the AH. Although no changes were observed in c-Fos expression in the AH, extensive prior work has found that this region contributes to the sex-specific regulation of aggression (Cheng et al., 2008; Delville et al., 1994; Ferris & Potegal, 1988; Terranova et al., 2016, 2017). 5-HT and AVP act in opposite ways within the AH to regulate aggression across sexes: AVP receptor activation promoted aggression in males but suppressed it in female Syrian hamsters, whereas 5-HT receptor activation decreased aggression in males and increased it in females (Terranova et al., 2016, 2017). Defining the phenotype of the Fos-positive neurons in the AH and other regions of the SBNN will be an important direction for future studies.

Finally, it is important to consider the broader implications of our findings. Sex differences in neuronal activity can be responsible for sex differences in behavior, but they can also function to ensure similar behavioral outputs can occur despite different underlying circuits (De Vries & Södersten, 2009). One possibility is that the differences we observed in neural activation enable males and females to display similar patterns of aggression toward same-sex conspecifics in competitive situations such as during the formation of dominance relationships, despite different hormonal environments. As noted previously, these differences may also contribute to higher levels of aggression observed in female hamsters compared to males (Ross et al., 2017). Another possibility to consider is that some of the differences observed may reflect other processes involved. For example, the sex differences we observed may mediate differences in conflict resolution and establishment of social hierarchy, which would affect how long the subjects engaged in aggressive behavior. Aggression is a key component of determining how animals engage with each other and form social hierarchies, impacting survival through competition for mates and resources. By identifying the mechanisms underlying aggression and how they differ between sexes, we gain insight into the broader organization of social behavior. Moreover, given that many neuropsychiatric disorders with prominent sex differences are also associated with maladaptive social skills, impulsive or reactive aggression, and agitation, understanding the neural basis of sex-specific aggression has significant translational relevance.

## Acknowledgements

Special thanks to Erica A. Cross for her assistance.

## Author Contributions: CRediT statement

**Susan D. Lee**: Conceptualization, Methodology, Formal analysis, Investigation, Writing – Original Draft, Visualization **Dario Aspesi:** Investigation, Validation, Review & Editing **Kim L. Huhman:** Methodology, Writing – Review & Editing **H. Elliott Albers:** Conceptualization, Resources, Writing – Review & Editing, Supervision, Funding acquisition

## Supplementary Figures

**Supplementary Figure 1.**
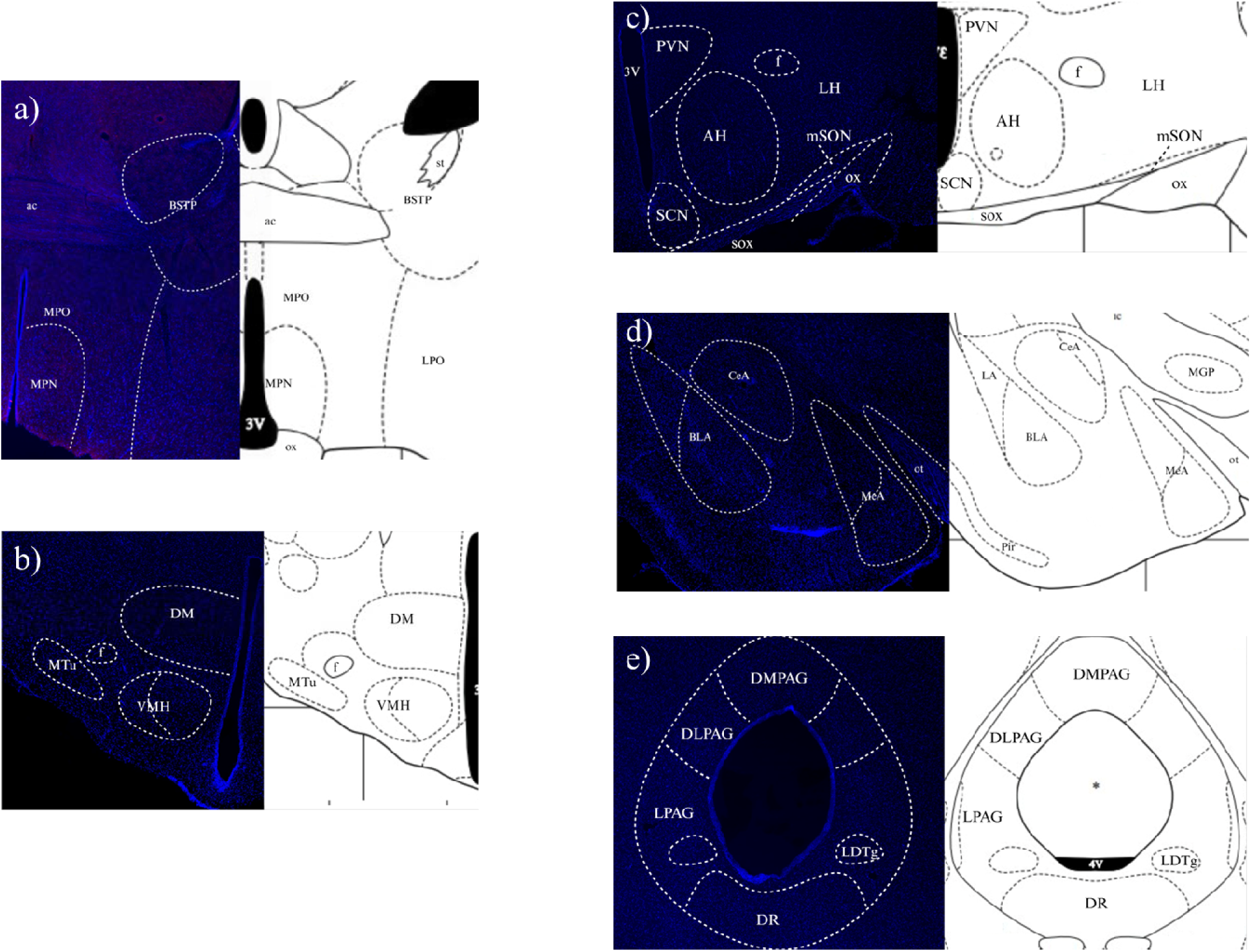
DAPI-stained sections of other regions of the SBNN, as defined by the Hamster Brain Atlas (Wood and Morin, 2001). a) Representative section of BNST and POA at 4X. BSTP = posterior BNST, MPO = medial preoptic area, MPN = medial preoptic nucleus, ac = anterior commissure. b) Representative section of VMH at 4X. DM = dorsomedial hypothalamic nucleus, f = fornix, Mtu = medial tuberal nucleus. c) Representative section of the PVN, AH, mSON, and LH. 3V = third ventricle, f = fornix, SCN = suprachiasmatic nucleus, sox = supraoptic decussation, ox = optic chiasm. d) Representative section of the amygdala at 4X. ot=optic tract, pir = piriform cortex, MGP= medial globus pallidus. e) Representative section of the DR and PAG (DMPAG, DLPAG, LPAG) at 4X. DMPAG = dorsomedial PAG, DLPAG = dorsolateral PAG, LPAG = lateral PAG, *=aqueduct, 4V = fourth ventricle, LDTg = lateral dorsal tegmental nucleus

